# Isolation and Molecular Identification of Endophytic Bacteria from *Tylosema esculentum*

**DOI:** 10.1101/2022.11.15.516524

**Authors:** Elisha Sinyinza, Percy M Chimwamurombe

**Affiliations:** Department of Biological Sciences, Faculty of Science, University of Namibia, Private Bag 13301, 340 Mandume Ndemufayo Avenue, Windhoek, Namibia; Department of Natural and Applied Sciences, Namibia University of Science and Technology, Private Bag 13388, Windhoek, 13 Jackson Kaujeua Street, Namibia, 9000.

**Keywords:** Bio-fertilisers, Endophytes, Nitrogen fixation, Symbiosis, Plant Growth-Promoting Bacteria

## Abstract

*Tylosema esculentum* is a perennial legume that is predominant in Namibia, Botswana, and South Africa. It is rich in nutrients, drought-tolerant, and a climate change contender for future agriculture. The objectives of this study were to isolate and identify culturable bacterial endophytes. Our study assessed the mean CFUs/mL among the tuberous roots, stems, and leaves to determine the density and diversity using ANOVA. It was a quantitatively controlled experiment in which seeds were grown in a greenhouse. The bacteria number varied from 1.29 × 10^8^ to 3.5 × 10^8^ CFU/ml of broth, with the highest in the tuberous roots and the lowest in the leaves. At a 5% significant level, we revealed that there is a difference among the bacterial densities of the plant parts. Shannon-Weiner diversity showed that there is a higher diversity of bacteria in the tuberous roots. Of the isolates that were obtained, 53.8% belonged to *Enterobacter*, 15.4% to *Bacillus*, 15.4% to *Pantoea*, and 7.67% to *Burkholderia*. These discoveries have made us more aware of the endophytic bacteria we now have. These strains might thus be used in food crops to boost productivity, enabling food security, especially in Africa. The significance of the isolates in biological control, agriculture, and biotechnology for growth promotion will be determined by further research.

## INTRODUCTION

*Tylosema esculentum* also known as marama bean and gemsbok, it is a long-lived underutilized perennial legume native to the arid region of Southern Africa.^4,10^ It belongs to the *Leguminosae* family and has three subfamilies namely, *Caesalpinioideae*, *Mimosoideae*, and *Palpilionoideae*.^4^ *T. esculentum* forms part of the indigenous people’s diet, and it is rich in protein and oil at about 29%–39% and 25%–28%, respectively.^10^

Endophytic bacteria colonize uninjured plant tissues without causing any harm to the plant, the interaction is mutualistic.^1,21^ Unfortunately, the relationship between endophytes and plants is not fully understood. The plant will provide nutrients for the bacteria and the bacteria will aid in the protection against pathogens, and secretion of gibberellins, phytopathogens, and cytokinins.^15^ Almost all, if not all plants are colonized by endophytic bacteria, some of which include nitrogen fixation and plant growth-promoting bacteria (PGPB). Endophytes normally reside in the intracellular spaces that contain a high amount of inorganic minerals, carbohydrates, and amino acids.^3,8^

The study of molecular biology has made strides that have increased our understanding of microorganisms and made it easier to identify different bacterial species by their molecular characteristics, which has resulted in the discovery of a vast array of bacteria. Due to the following distinctive characteristics, 16S ribosomal RNA (rRNA) is utilized to identify microorganisms. Because (I) it has a distinctive signature that is shared by all bacterial species, (II) it includes roughly 1500 base pairs (bp), which is a considerable amount of data, and (III) genes are quite accurate and do not vary over time. ^20^ We would then be able to recognize the endophytic bacteria that were present in *T. esculentum*. In addition, template bacterial DNA is frequently amplified using universal primers like 27F and 1492R.

In Southern Africa, *T. esculentum* tubers are employed as traditional medicines. ^4^ Rotaviruses are the main cause of diarrhoea in young children and persons with impaired immune systems. Marama leaf extracts have been shown to be effective microbicides against rotavirus. ^4,10^

Endophytic bacteria’s microbiome serves as a biocontrol agent and promotes growth, yield, and health. ^24^ These positive effects are characterised by and result from metabolic interactions. ^5^ Plant microsymbionts produce a wide variety of metabolites, which play a part in the interaction, competition, defense, and communication. ^24^ Endophytes synthesise necessary metabolites. ^5,9^ Endophytic bacteria have shown the potential in eliminating soil toxins by speeding up the phytoremediation process. They create several natural compounds that are beneficial to the hosts. ^6^ They are also known to serve as natural fertilisers by synthesising plant hormones like auxin, solubilizing phosphates, and fixing atmospheric nitrogen. ^12^

There is little knowledge of the endophytic bacteria that inhabit *T. esculentum’s* tissues. This research’s objective was to isolate and identify culturable endophytic bacteria. The results will increase our understanding of the many bacterial species that are there, which will be useful in figuring out how important the isolates are for biological control, agriculture, and biotechnology for growth promotion.

## MATERIALS AND METHODS

### Study design

It was a quantitatively controlled experimental study, and the major variables were endophytic bacteria and plant parts. We focused on the leaves (L), stems (S), and tuberous roots (TR). We had twelve (12) plants, and from each, three (3) parts were obtained, producing 36 replicates.

### Study site and sample collection

Soil samples were randomly collected from the University of Namibia at coordinates 22.6122° S, 17.0584° E. The soil was sterilised using an autoclave at 121 °C for 15 minutes. The soil samples were allowed to cool down for 3 hours prior to planting. The seeds were planted in 12 nursery pots and carefully placed on a flat surface in the greenhouse.

### Surface sterilisation

The fresh and healthy plant samples were washed thoroughly to remove sand particles. Surface sterilization was performed using 70% ethanol for 3 minutes, Sodium hypochlorite for 3 minutes, 70 % ethanol again for 30 seconds, and lastly washed them using distilled water.^9,11^ The disinfection process was confirmed by plating the aliquots of the final rinse in Tryptone Soy agar (TSA) at 30°C for 14 days.^8^

### Preparation of media

The modified SM-medium composed of 1g Malic acid, 1g Anhydrous glucose, and 20mg yeast extract in 1 liter (L) distilled water, was sterilized at 121°C for 15 minutes. This was followed by the addition of 1 ml vitamin stock solution (D-biotin 0.2g, p-Amino benzoic acid 0.02g, pyridoxine hydroxide 0.04g, thiamine dichloride 0.004g, riboflavin 0.02, nicotinic acid 0.04 plus 0.676 ml of distilled water) and 90% ethanol through a sterile syringe and a WHATMAN 0.2μm PES w/GMF filter by filtration. VM-ethanol was composed of the following ingredients per liter: 0.6g K_2_HPO_4_,0.4g KH_2_PO_4_, 0.5g NH_4_Cl, 0.2g MgSO_4_.7H_2_O, 1.1g NaCl; 0.026g CaCl_2_.2H_2_O; 0.01g MnSO_4_.H_2_O; 0.002g Na_2_MoO_4_.2H_2_O; 0.066g Fe (III)EDTA; 1g Yeast extract; 3g Tryptone; 12g bacteriological agar. The pH was adjusted to 7.56 using 0.5% sodium hydroxide and HCl. And then, the addition of 6ml of 90% ethanol and vitamin stock solution was done through filtration.^9,14^

### Endophytic bacteria isolation and culture

A sterile mortar and pestle were used to slice and macerate the plant samples, each weighing around 2 grams. The macerated materials were inoculated in SM broth devoid of nitrogen and left to incubate for 12 days at room temperature. Two weeks at 30 °C were followed by an incubation period at room temperature. ^4,8,10^

### Bacterial enumeration and pure culture

On VM ethanol plates, the tissue extracts from the SM broth were serially diluted from 10^−1^ to 10^−6^ times. After fully mixing the dilution tubes, duplicate aliquots of 0.1 ml of the dilution were inoculated on the VM-ethanol plates, and then disseminated using an L-shaped spreader. ^6^ The plates of VM-ethanol were incubated. The bacterial cell counts on the plates with 30 to 300 colonies were expressed as CFU per milliliter of liquid broth. Calculating it involved dividing the volume on the culture plate by the number of colonies multiplied by the dilution factor (0.1 ml). To facilitate processing and data analysis, the CFU/mL number was subsequently transformed into a logarithmic value. ^4,6,9^

Single colonies were sub-cultured using the four-quadrant streak technique before being incubated for seven days at 30°. For 48 hours, the pure colonies were inoculated in nutritional broth. The plates were kept in a 40 °C refrigerator. ^4^

### Genomic DNA extraction and Polymerase Chain Reaction

The bacterial cells were then centrifuged for 10 minutes at 10,000 rpm after being rinsed with 100 L of Ringer solution. The manufacturer’s instructions were followed while extracting DNA using ZR Fungal/Bacterial DNA MiniPrep™. Following the confirmation of the existence of DNA by the NanoDrop 2000, the DNA was placed onto 1% agarose stained with SYBR® Green I nucleic acid gel dye in 1X TBE buffer and examined under ultraviolet (UV) light to determine whether the quality and concentration are suitable for PCR. ^4^

The amplification of 16S rDNA was performed in a reaction with a final volume of 25 l containing 5 l of template DNA, 0.8 M of each of the primers 27F and 1492R, 5 l of nuclease-free water, and 12.5 l of GoTaq® Green Master Mix: Taq DNA polymerase, dNTPs, MgCl_2_, and buffers (Inqaba Biotechnology Industries, South Africa). A positive control (Bacteriophage lambda DNA with PCR mixture) and a negative control (PCR mixture without template DNA) were added to all the reactions. ^7^ The PCR profiles on the thermal cycler were as follows: Pre-denaturation (1 cycle) at 95 °C for 4 minutes; 35 cycles of denaturation at 95 °C for 30 seconds; annealing at 50 °C for 1 minute; 72 °C extension for 1 minute; and final extension (1 cycle) at 72 °C for 10 minutes; samples were stored at 4 °C. 7 The amplified DNA was observed on 1% agarose (100 ml of 1XTBE, 1 g of agarose, and 5 l of ethidium bromide) at 120 volts (v) for 35 minutes. The PCR products were sequenced and purified by Inqaba Biotechnical Industries (Pty) Ltd. in Pretoria, South Africa, using an ABI 3500XL sequencer. ^1,7^

### Molecular identification

Nucleotide sequences were edited and cleaned manually using Bioedit. The forward and reverse sequences were aligned using the complements of their sequences. All the sequences that were overlapping were trimmed to a consensus sequence that was generated using the sequence alignment editor. These consensus sequences were pasted in the blast search on the NCBI website (http://blast.ncbi.nlm.nih.gov/Blast.cgi).^9^

### Analyses of data

The findings of a normality test were applied to the mean CFU/mL values. At a 5% significance level, an ANOVA was carried out using SPSS version 23 to see if there is a difference in the bacterial density between the components. ^10^ The Shannon-Weiner diversity index on Microsoft Office 2013 (Excel), presuming that the species were repeated and randomly collected, was used to compare the bacterial diversity among the plant components. ^22,33^

## Results and Discussion

### Bacterial enumeration

The tuberous roots had the greatest CFU per ml average value, according to the mean CFU per ml measurements taken from the liquid broth of *T. esculentum* (Fig. 1). Because they are full of chemical substances, roots offer unique settings to a variety of microorganisms. ^3,6^ The soil microbiome and the bacterial niches in the root and rhizosphere have been revealed to be unique from one another. ^5,11^

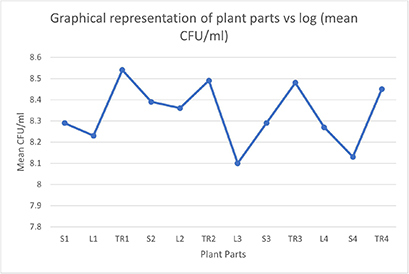

Root-based bacteria have an impact on a plant’s health and output. ^6^ It has been found that the behaviour of the root bacteria varies depending on which plant host they are associated with. ^9^

They can also change the interaction activity inside a single host because of changes in the host’s health or the environment. ^4,9,23^ Despite the significance of root microorganisms for plants and ecosystems, our knowledge of their organisational structure is relatively limited. This is a result of molecular biology’s glacial progress. Their rich diversity makes them challenging to research. There is evidence that the makeup of bacteria is influenced by both biotic and abiotic influences.

### Analyses of data

The Shapiro-Wilk test revealed that the CFU/ml of the bacterial samples were approximately normally distributed (p > 0.05), and normal Q-Q plots, histograms, and box plots were examined. 22 had a kurtosis of −0.064 (standard error = 1.22) and a skewness of 0.765 (standard error = 0.637). 10,22,23

### Analysis of Variances

The p-value (sig=0.013) in table 1 is less than 0.05. Thus, we find that there is enough data to support the argument that the bacterial densities among tuberous roots, stems, and leaves are different and reject the null hypothesis.

|  | Sum of<br>squares | DF | Mean<br>Squares | F | Sig. |
| --- | --- | --- | --- | --- | --- |
| **Between**<br>**groups** | 14.500 | 10 | 1.450 | 2.900 | 0.013 |
| **Within groups** | 0.500 | 1 | 0.500 |  |  |
| **Total** | 15.000 | 11 |  |  |  |

### Diversity indices

The maximum variety of endophytic bacteria was found in the tuberous roots (H value = 2.95), followed by the stems (H value = 1.09) and the leaves (H value = 0.563). (Fig. 2). The data show that there are variations in bacterial densities among the plant components. We may say that the bacterial density depends on the niche environment, and the bacterial composition depends on the materials present in a particular plant area as well as exposure to environmental elements like soil and sunshine.^15^ The Shannon-Weiner diversity index demonstrated that the tuberous roots had the highest endophytic bacterial diversity. Because practically all bacterial strains are present in the tuberous root, we may argue that it is the perfect plant component for isolating the necessary microorganisms.

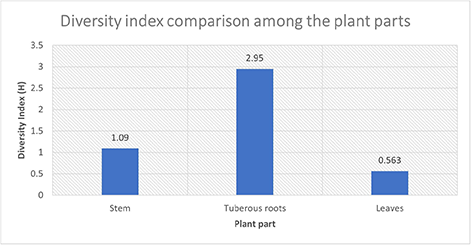

### Genomic DNA isolation and Polymerase Chain Reaction

The DNA was successfully isolated and amplified, and the results were confirmed by nanodroplet and gel electrophoresis (Fig. 3).

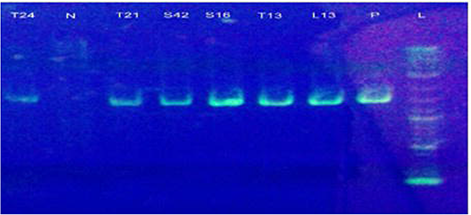

### Molecular Identification of Endophytic Bacteria

On the basis of the morphological characteristics of the colonies on the plates, we isolated 19 different strains of endophytic bacteria from *T. esculentum*. Thirteen isolates in all were successfully amplified. The endophytic bacteria were divided into the genera *Bacillus*, *Enterobacter*, *Pantoea*, and *Burkholderia* by bioinformatics analysis (Table. 2)

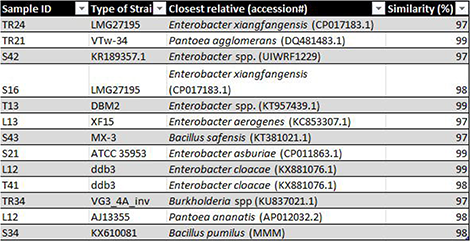

| Sample ID | Type of Strain | Closest relative (accession#) | Similarity (%) |
| --- | --- | --- | --- |
| TR24 | LMG27195 | *Enterobacter xiangfangensis* (CP017183.1) | 97 |
| TR21 | VTw-34 | *Pantoea agglomerans* (DQ481483.1) | 99 |
| S42 | KR189357.1 | *Enterobacter* spp. (UIWRF1229) | 97 |
| S16 | LMG27195 | *Enterobacter xiangfangensis* (CP017183.1) | 98 |
| T13 | DBM2 | *Enterobacter* spp. (KT957439.1) | 99 |
| L13 | XF15 | *Enterobacter aerogenes* (KC853307.1) | 97 |
| S43 | MX-3 | *Bacillus safensis* (KT381021.1) | 97 |
| S21 | ATCC 35953 | *Enterobacter asburiae* (CP011863.1) | 99 |
| L12 | ddb3 | *Enterobacter cloacae* (KX881076.1) | 99 |
| T41 | ddb3 | *Enterobacter cloacae* (KX881076.1) | 98 |
| TR34 | VG3_4A_inv | *Burkholderia* spp (KU837021.1) | 97 |
| L12 | AJ13355 | *Pantoea ananatis* (AP012032.2) | 98 |
| S34 | KX610081 | *Bacillus pumilus* (MMM) | 98 |

The most common genus, *Enterobacter*, made up 53.8% of the total, followed by *Bacillus* and *Pantoea* (15.4%) and *Burkholderia* (7.67%). Our research showed that the three species we recovered were also present in the sugarcane, rambutan, and *curcuma longa* fruit communities (*Nephelium lappaceum*). ^9,12,14^

The roots, stems, and leaves of sugarcane have all been discovered to contain *Enterobacter cloacae*. The bacterium is being utilised as a seed supplement before sowing as well as a post-harvest biocontrol for fungi that cause fungal diseases of fruits and vegetables. ^13^ Their findings proved that *E. cloacae* had characteristics of a plant growth-promoting bacterium (PGPB). ^6,8,17^ Clearly, the *E. cloacae* strain might be employed to promote the development of *T. esculentum* and other potential food crops.

It had been discovered that *Enterobacter aerogenes* produces acetoin, which induces resistance against phytopathogens. ^13, 20^. Rhizobacteria produce metabolites, which are the intermediaries in the process. ^13,15,16^ *E. aerogenes’* production of acetoin has helped maize plants develop efficient defences against the phytopathogen *Setosphaeria turcica*. ^13,17^ We detected *E. aerogenes* from the tuberous roots, and it was previously found to coexist with the root tissues. The results support those from prior studies. ^6,14,26^ It is clear that *E. aerogenes* is capable of fixing atmospheric nitrogen and offering protection from phytopathogens.

*Enterobacter asburiae* is a gram-negative, non-spore-forming rod-shaped bacterium from the *Enterobacteriaceae* family. ^24^ It lives on plants, soil, and water surfaces. ^15^ Most strains of *E. asburiae* can inhibit a variety of plant diseases by protecting roots, stems, leaves, and seeds from fungal infection. ^15, 22^. It has been discovered that *E. asburiae* has the ability to promote plant growth, auxin production, and nitrogen fixation in the atmosphere. ^15,16^

Plants, water, and soil are just a few of the several settings where *Enterobacter xiangfangensis* can be found. It grows best at 30 °C. ^17^ IAA (Indole-3-acetic acid), which is necessary for cell development, elongation, and the solubilization of phosphates, and can be synthesised by *E. xiangfangensis*. ^5,16,17^

*Enterobacter* spp. promotes plant development. It lives in the soil but is mostly found in plant tissues where it has a mutualistic interaction. ^21,21^ It has been described as a nodule-forming, nitrogen-fixing bacterium. ^8^ Recent research has demonstrated that the *Enterobacter* spp. produces 2,3-butanediol, IAA, siderophores, and ACC (1-aminocyclopropane-1-carboxylate deaminase), all of which are known to promote plant development. ^12^ Bacteria with features that promote plant development are frequently found in the environment. ^18^

Previously known as *Enterobacter agglomerans* or *Erwinia herbicola*, *Pantoea agglomerans* is a diazotrophic, non-spore-forming, rod-shaped endophytic bacteria. ^3,14^ It has frequently been isolated from seeds, mandarin oranges, fruits, plants, and animal or human excrement. ^19^ In the range between 25 and 30 °C, the bacteria may thrive. It is also known to thrive in settings with a lot of salt or little water. ^15^ Additionally, *P. agglomerans* has been discovered to be a bacterium that promotes plant development. ^4^

A member of the *Enterobacteriaceae* family, *Pantoea ananatis* is characterised by its ubiquity and is connected to both plant and animal hosts. *P. ananatis* is a member of the endophytic and epiphytic flora and has been isolated from the roots, leaves, and stems of a variety of plants. It has been discovered on banana leaves and rice seeds as an endophyte. ^10, 22^ *P. ananatis* pathogenic strains that infect plants have been discovered in other studies. ^6^ It has been discovered that secondary metabolites isolated from *P. ananatis* have anti-fungal properties that are effective against *Mycosphaerella musicola*. ^22^

*Burkholderia* spp. are adaptable bacteria that may live in a variety of habitats. The bacteria grow slowly. ^5,6,24^ *Burkholderia* spp. is mostly employed as a biocontrol agent, for bioremediation, to prevent bacterial or fungal plant infections, and to enhance plant development. ^23^ *Burkholderia* spp. were found in China’s tobacco rhizosphere in recent research. ^23^ According to reports, it creates antimicrobial compounds, which can provide scientists valuable insights into the synthesis of bio-control agents and the elimination of plant diseases. Additionally, *Burkholderia* spp. isolated from rice has traits that support plant development, including the production of ACC deaminase, the capacity to fix nitrogen, and antifungal properties. ^7^

Although it may also be found in other environments including water and soil, *Bacillus safensis* is mostly found as an endophyte in plants. ^23^ To survive in harsh conditions, *B. safensis* depends on its genotypic and physiological characteristics. ^23^ According to research, *B. safensis* possesses characteristics that encourage plant development and hold potential for use in biotechnology. As a biocontrol agent. ^7,21^

By using resistance and direct antagonistic interactions, the endophytic bacteria *Bacillus pumilus* has been exploited as a biocontrol agent against other plant diseases and soil-borne pathogens. ^17^ In this investigation, a stem-isolated *B. pumilus* was used. It has also been discovered in a sick cucumber plant’s stem. ^16^ It is known that bacteria may create resistance in plants to a variety of diseases. ^21^

We were able to effectively isolate 12 distinct bacterial endophytes from *T. esculentum’s* tissues. We can certainly state that the isolates we acquired are nearly identical to those in the database since the isolated endophytes detected had an excellent homology of 97% to 99%. Our results are in line with other studies; the endophytic bacteria we found are well-known endophytes seen in grasses, sugarcane, and other plants. Endophytic bacteria are members of a class of plant growth-promoting bacteria that also generate the hormones auxin and gibberellin, which are thought to have anti-pathogenic properties.

These bacteria also assist plants in protecting themselves from bacterial and fungal infections, metals, organic pollutants, salt, floods, droughts, and flower withering in addition to directly boosting plant development. Future research will focus on the characterisation of endophytic bacteria and how they may be used to biocontrol bacterial and fungal diseases that affect this important legume for human consumption.

